# Cross-tissue transcriptome-wide association study prioritizes druggable targets for hepatocellular carcinoma

**DOI:** 10.64898/2025.12.18.695261

**Authors:** Jiahuizi Peng, Wen Zhang

## Abstract

**Background:** Hepatocellular carcinoma (HCC) is known as the sixth most common cancer and the third leading cause of death related to cancer. While genome-wide association study (GWAS) has uncovered risk loci, mapping variants to causal genes and tissues is challenging. The aim of this study is to translate HCC GWAS signals from variants to genes and tissues to identify genetically regulated causal candidates and potential therapeutic targets.

**Methods:** We collected and integrated 12 benchmark GWAS summary statistics from different research including various areas and causes of HCC. We applied S-PrediXcan using GTEx 49 tissues to infer genetically regulated expression. Multiple testing was controlled using group-wise Benjamini-Hochberg FDR within tissues (primary, q≤0.05). For visualization, suggestive signals with p≤0.01 were highlighted; heatmaps display FDR-thresholder results, whereas other figures show the full distribution. We then performed cross-tissue prioritization, gene-trait network construction, gene set enrichment analysis (GSEA), and pathway enrichment.

**Results:** We identified 422 significant gene–tissue associations (BH-FDR within tissue, q≤ 0.05) regarding HCC, consolidating to 34 unique genes. Top tissues included breast mammary, left ventricle, and esophagus muscularis. Excluding established HCC loci (e.g., PNPLA3, TM6SF2, TERT, HSD17B13), we prioritized 32 druggable “novel” candidates, with top-ranked examples including DHCR24 and HLA-DPB1.

**Conclusions:** Cross-tissue TWAS integrating GWAS and GTEx models identifies genetically regulated genes associated with HCC, refines tissue specificity, and prioritizes plausible drug targets. This gene-centric framework complements variant-level GWAS and supports mechanism-guided prevention and therapy development.

## Introduction

Hepatocellular carcinoma (HCC) continues to rise in both incidence and mortality, driven by the growth of chronic liver disease and cirrhosis, challenges in early detection, population aging, and high post-treatment recurrence [1]. Although HCC is relatively uncommon in the United States, it is prevalent in sub-Saharan Africa and Southeast Asia, disproportionately affecting male over 50 years of age [2]. Major risk factors include chronic hepatitis B and C infections, alcohol-related liver disease, and cirrhosis [3–4]. This epidemiologic and etiologic heterogeneity underscores the need to delineate the genetic and tissue contexts through which risk manifests.

Genetic susceptibility contributes substantially to HCC risk, and recent genome-wide association studies (GWAS) have identified multiple loci across etiologies and populations [5]. For instance, non-viral HCC susceptibility loci in European populations include PNPLA3, TM6SF2, and TERT; additional signals such as KLF15, HSD17B13, APOE, and HFE have been reported across large multi-cohort analyses; variants at WNT3A-WNT9A, TM6SF2, and PNPLA3 predispose individuals with alcohol-related liver disease to HCC; and PNPLA3 and SAMM50 risk alleles increase HCC susceptibility in U.S. cohorts [6–8]. While these findings highlight the genetic architecture of HCC and inform risk stratification, most variant-level associations do not directly reveal the causal genes, tissues, or mechanisms — limiting translation to biomarkers and therapeutic targets [9].

Mounting evidence shows that disease-associated variants are enriched in cis-regulatory elements (promoters, enhancers) and frequently modulate gene expression. This has motivated transcriptome-wide association studies (TWAS), which integrate GWAS summary statistics with gene expression prediction models to test gene-level associations with complex traits [10]. MetaXcan (including S-PrediXcan and S-MultiXcan) [11–12] is a widely adopted transcriptome-informed framework that leverages Genotype-Tissue Expression (GTEx) models from PredictDB to infer genetically regulated expression and its relationship to disease across tissues. By aggregating the combined effects of multiple cis-SNPs, TWAS approaches improve interpretability, refine tissue-of-action, and enable downstream functional prioritization — spanning candidate gene validation, pathway analysis, and drug target nomination [13].

Here, we systematically integrate HCC GWAS with GTEx-based S-PrediXcan models to translate SNP-level signals into gene- and tissue-level insights [14]. By combining multiple HCC GWAS spanning diverse populations and etiologies with 49 tissue-specific prediction models, we aim to: (i) identify genetically regulated genes associated with HCC; (ii) refine tissue specificity to illuminate liver – immune – metabolic axes of risk; and (iii) prioritize plausible therapeutic targets supported by gene-level evidence, tissue context, and pathway coherence. This gene-centric framework complements variant-level GWAS, advances mechanism-guided risk stratification, and provides a foundation for functional studies and translational development in HCC.

## Methods

### Study design and overview

We performed a multi-cohort, multi-tissue transcriptome-wide association study (TWAS) to translate hepatocellular carcinoma (HCC) GWAS signals from SNPs to genes and tissues. We harmonized 12 independent GWAS summary datasets, applied S-PrediXcan with GTEx v8 prediction models across 49 tissues [15], controlled multiple testing using group-wise Benjamini-Hochberg false discovery rate (BH-FDR) within tissues (primary, q≤0.05) [16], and conducted cross-tissue prioritization, gene–trait network analysis, and pathway enrichment. Heatmaps display FDR-thresholder results; other figures show full distributions with suggestive signals highlighted at p≤0.01.

### GWAS data sources and inclusion

We curated summary-level GWAS for hepatocellular carcinoma (HCC) and closely related liver traits, uniformly screening inputs for required fields prior to downstream processing. Included datasets were drawn from the GWAS Catalog (GCST90043858, GCST90041897) and OpenGWAS (finn-b-C3_LIVER_INTRAHEPATIC_BILE_DUCTS_EXALLC, ieu-b-4915, finn-b-CD2_BENIGN_LIVE_BILE_EXALLC, finn-b-CD2_BENIGN_LIVER_EXALLC, bbj-a-158, ebi-a-GCST90018858, ebi-a-GCST90018638, ebi-a-GCST90018583, ebi-a-GCST90018803, ebi-a-GCST90092003). Eligibility required summary statistics reporting at minimum effect allele, non-effect allele, effect size or z-score, standard error, and p-value. Details of all GWAS datasets, including accession IDs, ancestry, phenotype definitions, and sample sizes, are provided in Supplementary Table 1. Cohorts with predominantly European or East Asian ancestry were prioritized, with additional ancestries included when available. When reported by source studies, we documented genomic inflation metrics and LDSC intercepts to contextualize confounding versus polygenicity; no recalibration of the supplied statistics was performed.

### Harmonization and preparation

We converted OpenGWAS VCF datasets to tabular formats (gzipped TSV) by extracting scalar statistics from the FORMAT column (ES, SE, LP, EZ). For each variant, we set effect_allele = ALT and non_effect_allele = REF; computed *p* = 10^−LP^ (falling back to INFO.PVAL if present and LP was missing); and set *z* = ES/SE when both were available with SE ≠ 0, otherwise *z* = EZ. The parser is compatible with single-sample VCFs and uses the first sample in multi-sample VCFs; variant_id preferred the native VCF ID and otherwise used chr: pos: ref: alt. All 12 GWAS were lifted to GRCh38 using hg19ToHg38 chain files and verified post-liftover via allele/position checks, then harmonized to the MetaXcan 1000 Genomes reference to standardize chromosome formats and positions and to derive panel_variant_id [17]. Large files were processed in chunks (for example, CHUNK_LINES = 1.5 × 10^6^, triggered at SPLIT_THRESHOLD = 3.0 × 10^6^) and merged under a unified header. Outputs followed a fixed order: variant_id, panel_variant_id, chromosome, position, effect_allele, non_effect_allele, p, z, effect_size, standard_error, sample_size, n_cases; when unavailable, sample_size=0 and n_cases=0 were inserted as placeholders. Allele alignment and quality control followed conservative rules: ambiguous palindromic SNPs (A/T, C/G) were removed or retained only when minor-allele frequency permitted strand inference; where available, SNPs with low imputation quality (e.g., INFO/RSQ < 0.8) or very low frequency (e.g., MAF ≤ 0.01) were excluded. Integrity checks included gzip validation, header/column consistency, and verification that *z* ≈ ES/SE where applicable. Original phenotype labels were normalized to canonical trait identifiers prior to analysis (Supplementary Table S2).

### TWAS analyses (S-PrediXcan; optional S-MultiXcan)

We performed transcriptome-wide association using S-PrediXcan on GTEx v8 (PredictDB) MASHR-based weights across 49 tissues [18], including liver (approximately *n* ≈ 208). Model-matched SNP covariance was used as provided by the MetaXcan resources. For each GWAS, S PrediXcan was run across all 49 tissues, generating 49×12 gene–tissue association files with *z*-scores and *p*-values. Cross-tissue evidence was summarized by aggregating single-tissue results (for example, counts, heatmaps, prioritization). When applied, S-MultiXcan integrated per-gene signals across tissues leveraging cross-tissue eQTL sharing. Multiple testing control used Benjamini-Hochberg FDR within tissues (primary, *q* ≤ 0.05). Across tissues and traits, the matrix of tested genes and grouped BH–FDR–significant associations (*q* ≤ 0.05) is provided in Supplementary Table S3. For exploratory visualization, suggestive signals with *p* ≤ 0.01 were highlighted. For conservative reporting, Bonferroni thresholds were defined as *α* = 0.05/*m*, where *m* is the number of effective tests and detailed in Supplementary Methods. Reporting policy: heatmaps display FDR-thresholder associations (BH within tissue, *q* ≤ 0.05); volcano plots for each GWAS display all tests with significance classes (FDR-significant versus suggestive *p* ≤ 0.01) highlighted.

### Tissue specificity and enrichment

For each tissue, we computed the number of significant gene-tissue associations (BH-FDR, *q* ≤ 0.05), total tested genes, and the proportion *N*_sig_/*N*_total_. Enrichment analyses, where applicable, used hypergeometric or permutation tests relative to each tissue’ s testable gene background, with multiple-testing correction by BH (*q* ≤ 0.05). Gene × tissue heatmaps were generated using −log_10_(*q*) for significant cells (others masked), with hierarchical clustering to reveal cross-tissue modules and axes (for example, liver-immune-metabolic).

### Gene–trait network (Cytoscape)

We constructed a bipartite network linking GWAS traits to genes via significant associations, retaining edges at BH FDR *q* ≤ 0.05. We summarized topology by node degree and betweenness centrality to identify hubs and detected communities using Louvain or MCODE [19]. Visualization settings encoded degree/centrality by node size and tissue/trait categories and effect directions by node/edge color where available. Networks were constructed from BH-FDR significant associations (*q* ≤ 0.05). Node and edge tables are provided in Supplementary Data S1–S4 together with an MHC-focused subset and node metrics.

### Pathway enrichment (GSEA)

We ranked genes by signed TWAS statistics using a unified scheme (−log _,-_(*p*)), with the background defined as the union of testable genes. Gene set libraries included MSigDB Hallmark, GO Biological Process, and Reactome. We used fgsea for enrichment testing and defined significance at BH FDR (q≤0.05) [20]. Redundant GO terms were simplified, and cross-tissue duplicates were summarized. We reported top terms per library and showed representative enrichment curves (for example, REACTOME_INTERFERON_GAMMA_SIGNALING).

### Druggability and translational prioritization

We derived a prioritization table (TargetScore) that integrates multiple sources of evidence and TWAS-significant gene sets. Inputs comprised summary metrics including but not limited to: G (maximum −log _,-_(*p*) across traits), R (recurrence across traits/tissues), S (pathway support aggregated from significant GSEA leading-edge genes), T (drug targetability proxy), E (optional external evidence penalization), and H (auxiliary heuristics). Filtering considered overlap with TWAS-significant genes [21]; high-priority candidates were defined by a pre-specified TargetScore threshold ( ≥0.7). Optionally, composite prioritization can integrate TWAS significance, tissue specificity, network centrality, colocalization, and pathway coherence. We reported the number of druggable candidates and highlighted top-ranked examples [22]. Known HCC risk genes (for example, PNPLA3, TM6SF2, TERT, HSD17B13) were flagged and excluded from the “novel” list [23].

### Visualization and figure policy

We derived a prioritization table (TargetScore) that integrates multiple sources of evidence and TWAS-significant gene sets. Inputs comprised summary metrics including Volcano plots were generated per GWAS using all tests, with significance classes highlighted (BH-FDR *q* ≤ 0.05; suggestive *p* ≤ 0.01). Gene × tissue heatmaps showed FDR-filtered significance using −log _,-_(*q*) [11]. Gene-trait networks were rendered in Cytoscape using width of edges to show significance [24]. A flowchart summarized the end-to-end pipeline from GWAS harmonization to TWAS, network analysis, GSEA, and prioritization.

### Software and versions

Analyses used MetaXcan (S-PrediXcan) with GTEx v8 (PredictDB; MASHR weights) [25]. We used R and Python environments with key R packages including dplyr, data.table, ggplot2, pheatmap, igraph, fgsea, readr, and Cytoscape for network visualization. Reproducibility materials (code and environment files, Github) will be made available; random seeds were fixed where applicable.

## Results

### Tissue specificity and enrichment

Within-tissue FDR proportions (significant pairs per total testable genes) ranked Breast_Mammary_Tissue, Heart_Left_Ventricle, and Esophagus_Muscularis among the top tissues. A matrix-based aggregation across traits (rowwise summation/normalization) similarly elevated extrahepatic tissues, including Brain_Spinal_cord_cervical_c-1, Brain_Nucleus_accumbens_basal_ganglia, and Heart_Left_Ventricle. While such rankings are unconventional for a liver cancer, they underscore systemic and extrahepatic contributions—metabolic, cardiovascular/stromal, and neuroimmune — to HCC risk and progression. In parallel, the gene × tissue heatmap (Figure 2) reveals a clear “liver/immune” dual axis: liver/hepatobiliary tissues and immune/blood tissues show clustered significance, aligning with enrichment of metabolic reprogramming and inflammation/antiviral response pathways. The column-wise z-standardized matrix of significant ratios (sig/all) used for heatmap visualization is available in Supplementary Table S4, with 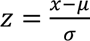 per trait.

**Figure 1.**
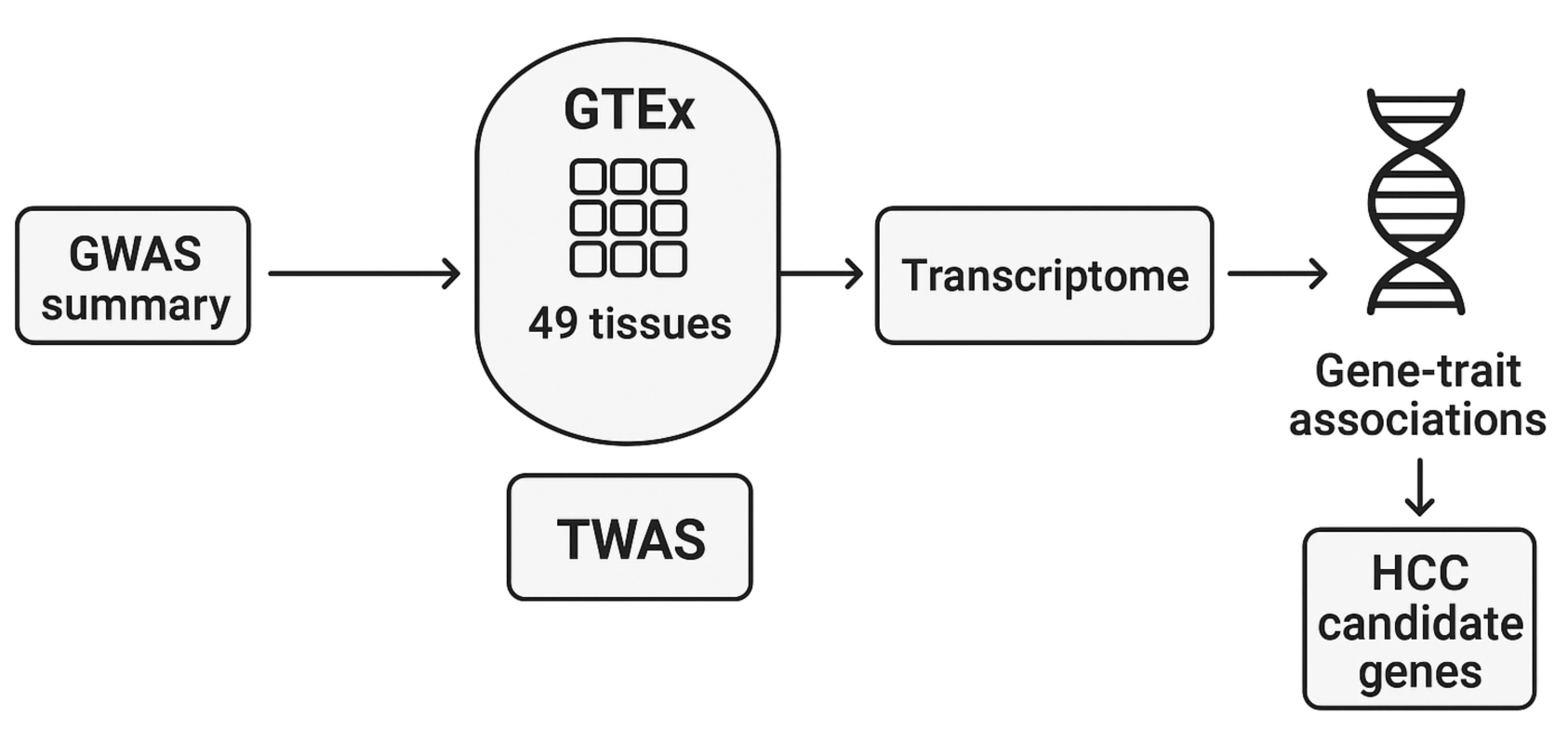
Variant-to-gene translation via cross-tissue TWAS. Summary statistics from 12 hepatocellular carcinoma (HCC) GWAS are harmonized (liftover to GRCh38, allele alignment, QC) and integrated with GTEx v8 expression prediction models across 49 tissues. Using S-PrediXcan, we test associations between genetically regulated expression and HCC across tissues, generating gene–trait association statistics. Significant associations are defined by within-tissue Benjamini-Hochberg FDR (primary *q* ≤ 0.05); suggestive signals with *p* ≤ 0.01 are flagged for visualization. Significant gene – trait pairs are consolidated into HCC candidate genes and carried forward to cross-tissue prioritization, gene – trait network construction, and pathway enrichment. Abbreviations: GTEx, Genotype-Tissue Expression; TWAS, transcriptome-wide association study; BH-FDR, Benjamini-Hochberg false discovery rate.

**Figure 2.**
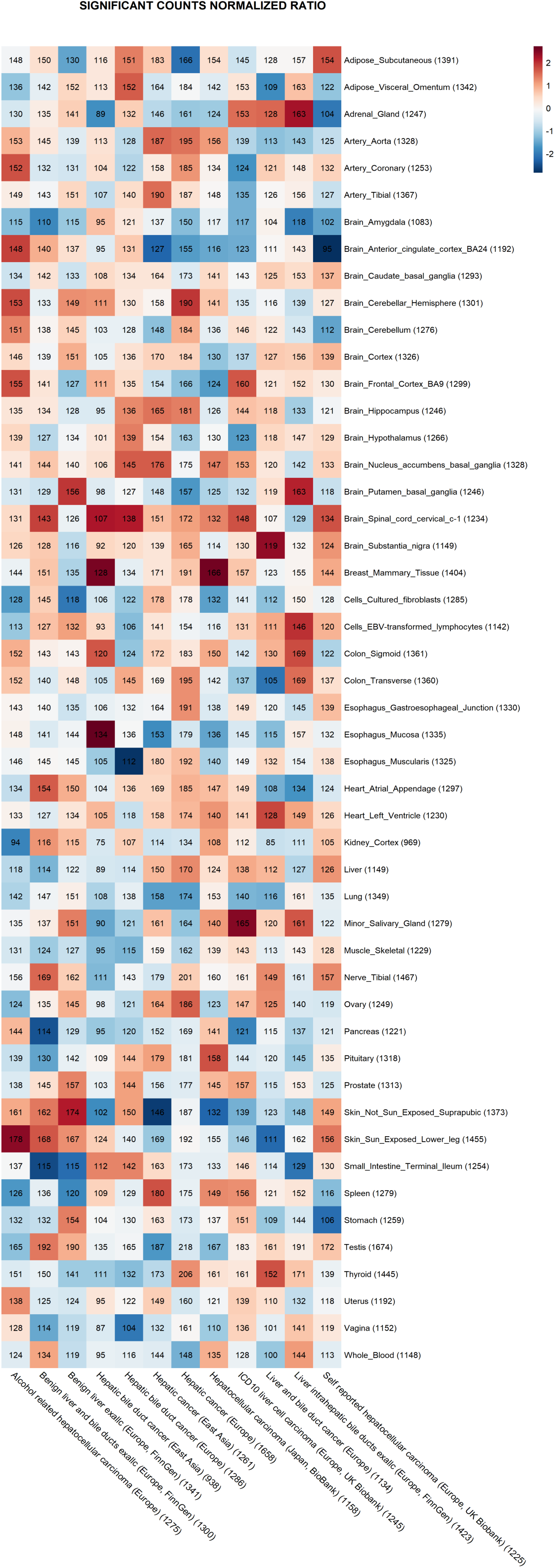
Gene × tissue heatmap of FDR-significant associations across 49 GTEx v8 tissues and 12 HCC-related GWAS traits. Each cell reports the raw number of significant genes (*N*_sig_, printed) for a given tissue-trait pair under within-tissue BH-FDR control (primary *q* ≤ 0.05). Tissue labels include, in parentheses, the number of testable genes in that GTEx model (*N*_total_; for example, Breast_Mammary_Tissue = 1404, Heart_Left_Ventricle = 1230, Esophagus_Muscularis = 1325, Liver = 1149, Whole_Blood = 1148). To enable cross-tissue comparison, we computed a normalized ratio *R* = *N*_sig_/*N*_total_ for each cell and then centered and scaled within tissue (row-wise z-score). The color scale encodes this normalized ratio (red = above-average within that tissue; blue = below-average; approximate range [−2, 2]). Rows (tissues) and columns (traits) appear in a fixed order. Abbreviations: BH-FDR, Benjamini-Hochberg false discovery rate; GTEx, Genotype-Tissue Expression; GWAS, genome-wide association study; HCC, hepatocellular carcinoma.

### Gene-trait network and hub genes

Using FDR-significant associations (edges retained at *q* ≤ 0.05), we built a bipartite GWAS/trait-gene network in Cytoscape. The network exhibits distinct communities corresponding to immune activation/antiviral response, lipid/cholesterol metabolism, and cell cycle/proliferation themes. Several genes emerge as hubs by degree and betweenness centrality, bridging multiple GWAS and tissue modules, highlighting potential regulators of HCC pathogenesis. Figure 3 annotates representative hubs and community assignments; full Cytoscape node/edge tables and MHC subsets are available as Supplementary Data SD1–SD4.

**Figure 3.**
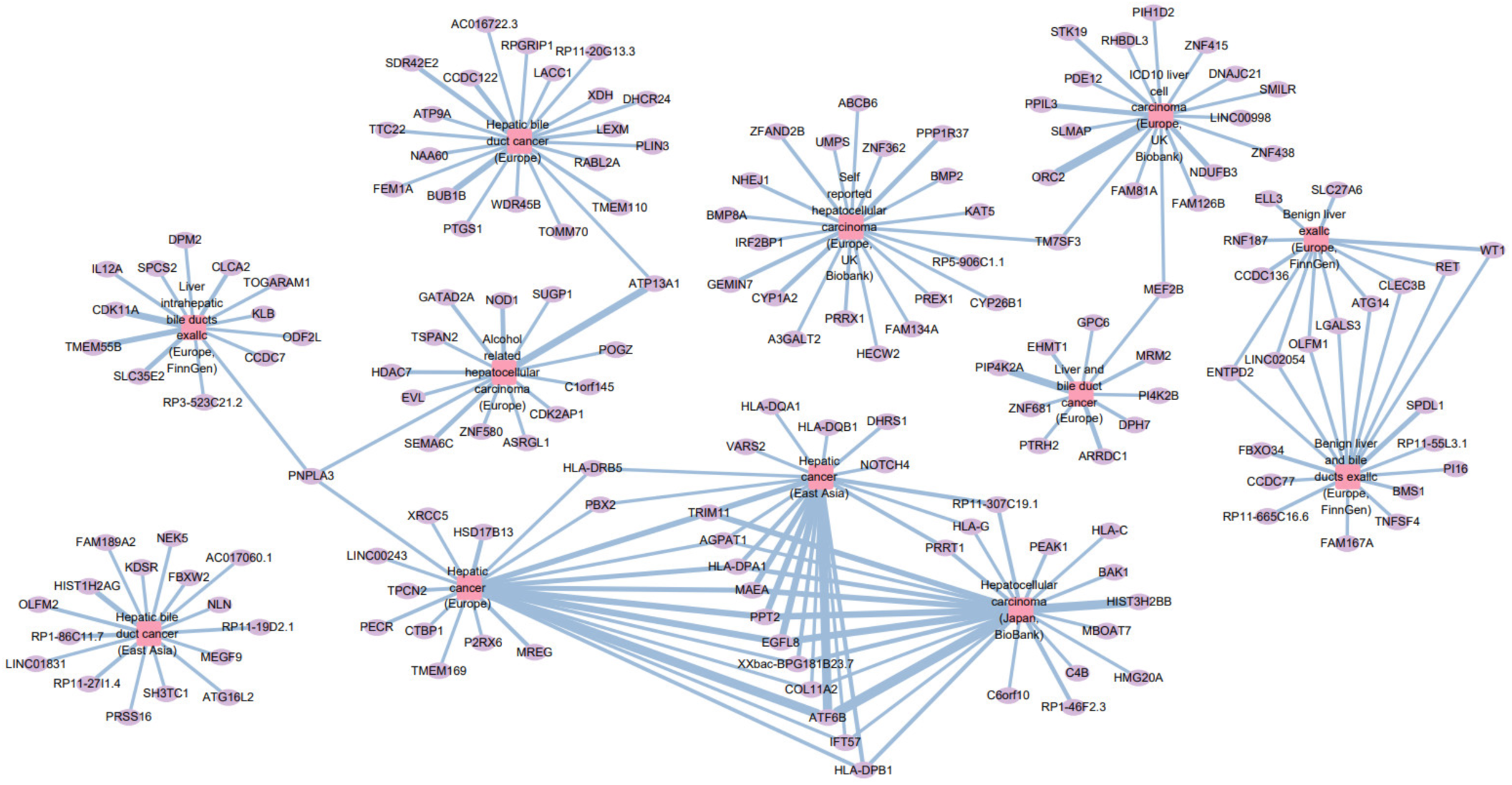
Bipartite gene–trait network of FDR-significant associations (*q* ≤ 0.05). Squares represent GWAS traits and circles represent genes; an edge denotes a significant gene–trait association in at least one tissue. Node size scales with centrality (degree/betweenness), and edge width reflects recurrence (e.g., number of GWAS or tissues supporting the edge). Community detection (Louvain/MCODE) reveals modules consistent with immune presentation/antiviral signaling (e.g., HLA class II cluster), lipid/cholesterol metabolism, and cell-cycle/proliferation. Hub genes bridging multiple traits/tissues are labeled. This network highlights cross-GWAS concordance and tissue-context organization of TWAS signals.

**Table1.**
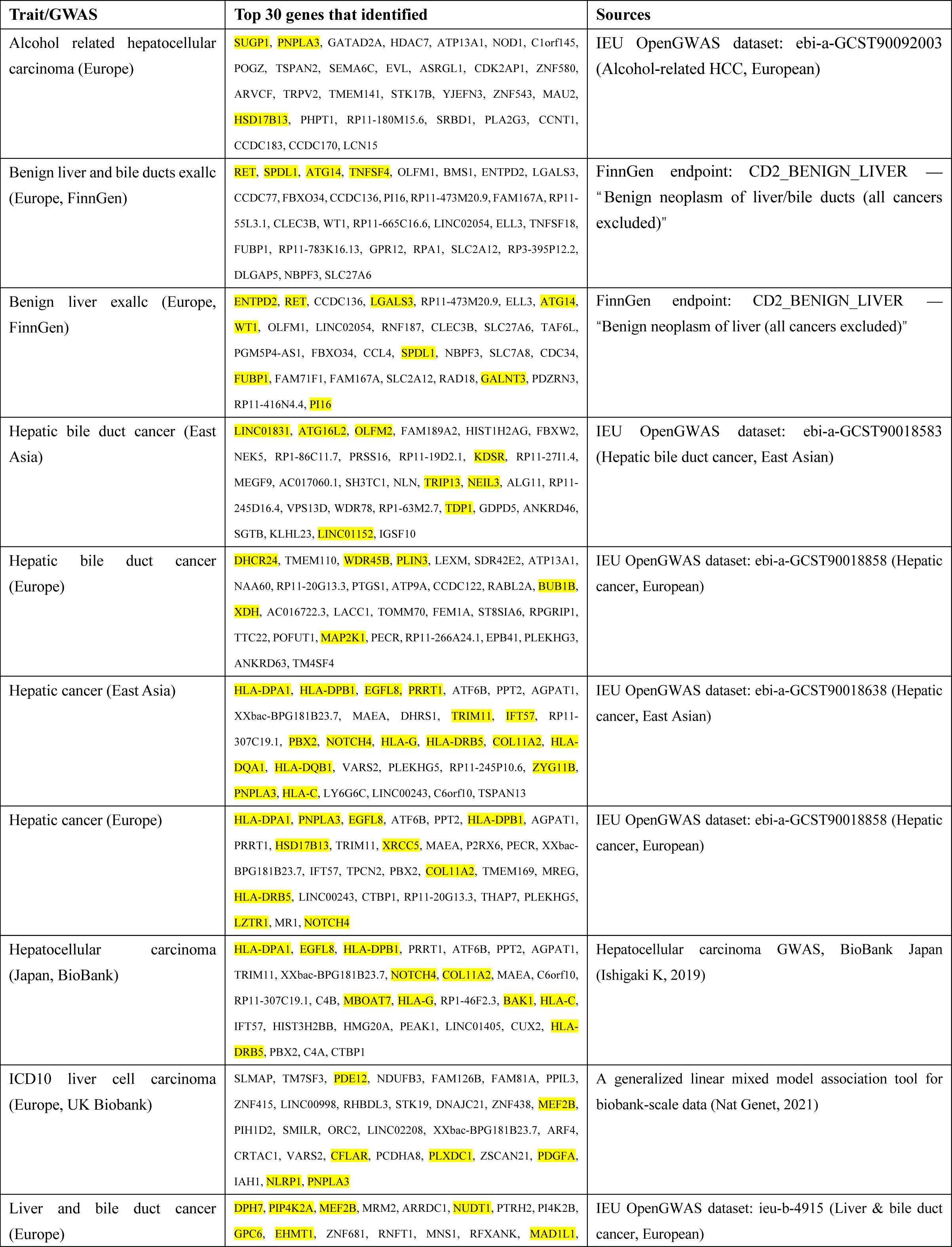
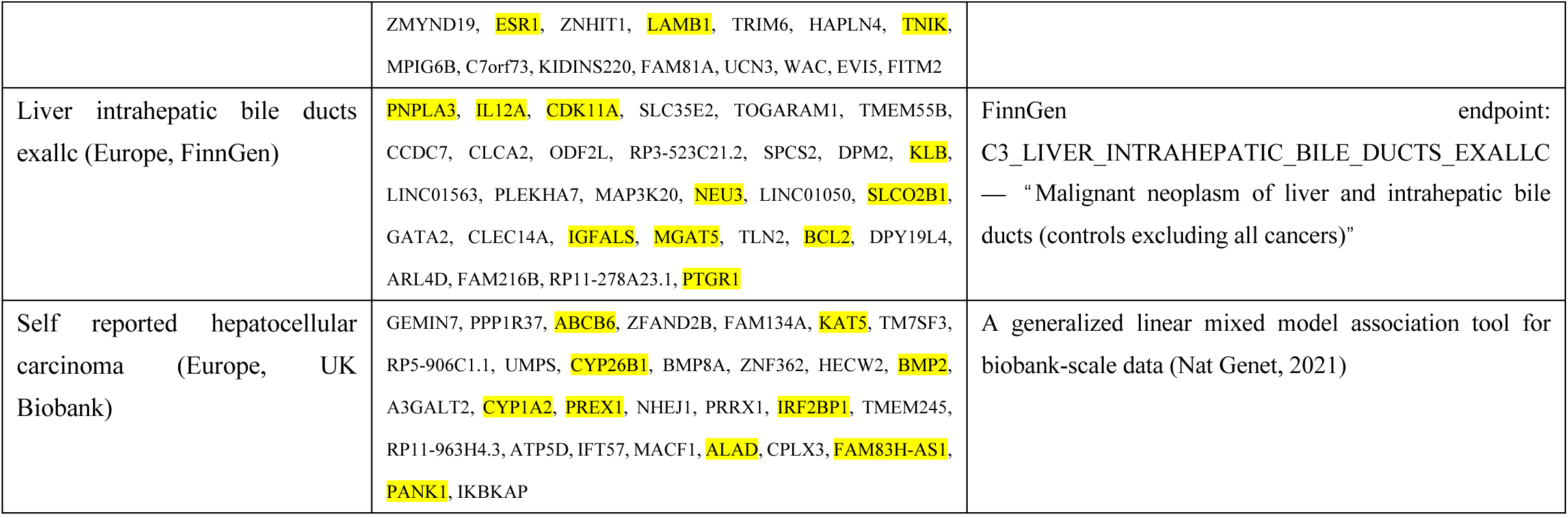
Top 30 unique TWAS genes per HCC-related GWAS trait aggregated across 49 GTEx v8 tissues. Highlighted genes have previously been verified as HCC-related genes.

### Consistency and heterogeneity across GWAS/etiologies

Per-tissue volcano panels for all 12 traits (48 tissues, excluding Liver) visualize TWAS z-scores versus −log _10_(*p*) with Top-20 labels (Supplementary Figure S1). Trait-level volcano overlays across 49 GTEx tissues (points colored by tissue) highlight trait-specific significance landscapes (Supplementary Figure S2). Per-tissue volcano panels for all 12 traits (48 tissues, excluding Liver) visualize TWAS z-scores versus −log _10_(*p*) with Top-20 labels (Supplementary Figure S1). Across hepatocellular carcinoma (HCC) and related liver-trait GWAS, we compiled the top 30 unique TWAS genes per trait (non-redundant within trait) by ranking per-tissue results and aggregating across 49 GTEx v8 tissues. For each canonical trait, the Top-30 genes (ranked by the maximum −log_10_(*p*) across tissues) are listed in Supplementary Table S5. Genes were prioritized by rank (primary), or by −log _10_(*p*) / ∣ z ∣ when rank was unavailable. Recurrent candidates appearing in multiple, independent GWAS form a coherent immune–metabolic theme: lipid droplet and neutral lipid handling (for example, PNPLA3, TM6SF2, HSD17B13), immune/HLA axis (for example, HLA-DPA1, HLA-DPB1; consistent with complement biology), and developmental/differentiation signaling (for example, NOTCH4). These gene-level recurrences align with our pathway findings (complement activation, interferon-γ signaling, triglyceride/neutral lipid metabolism), supporting an immune–metabolic coupling narrative for HCC susceptibility. We quantified across-GWAS replication as *R_g_* (the number of distinct GWAS in which gene *g* appears) and flagged cross-ancestry support when a gene recurred in both European and East Asian cohorts. Known HCC risk genes (PNPLA3, TM6SF2, HSD17B13) were labeled and excluded from the “novel” list; the remaining recurrent genes, especially those concordant with enriched pathways, constitute higher-priority candidates for follow-up. Full replication statistics (*R_g_*), ancestry coverage, and theme labels are provided in Supplementary Tables.

### Pathway enrichment and biological mechanisms (Figure 6)

We performed targeted gene set enrichment analysis (GSEA) across MSigDB Hallmark, GO Biological Process (GO BP), and Reactome libraries using genes ranked by signed TWAS statistics, with pathway significance defined at q ≤ 0.05 (Benjamini–Hochberg) [26]. A concise set of significant pathways was identified across libraries (Hallmark, n=1; GO BP, n=3; Reactome, n=1).

**Figure 4.**
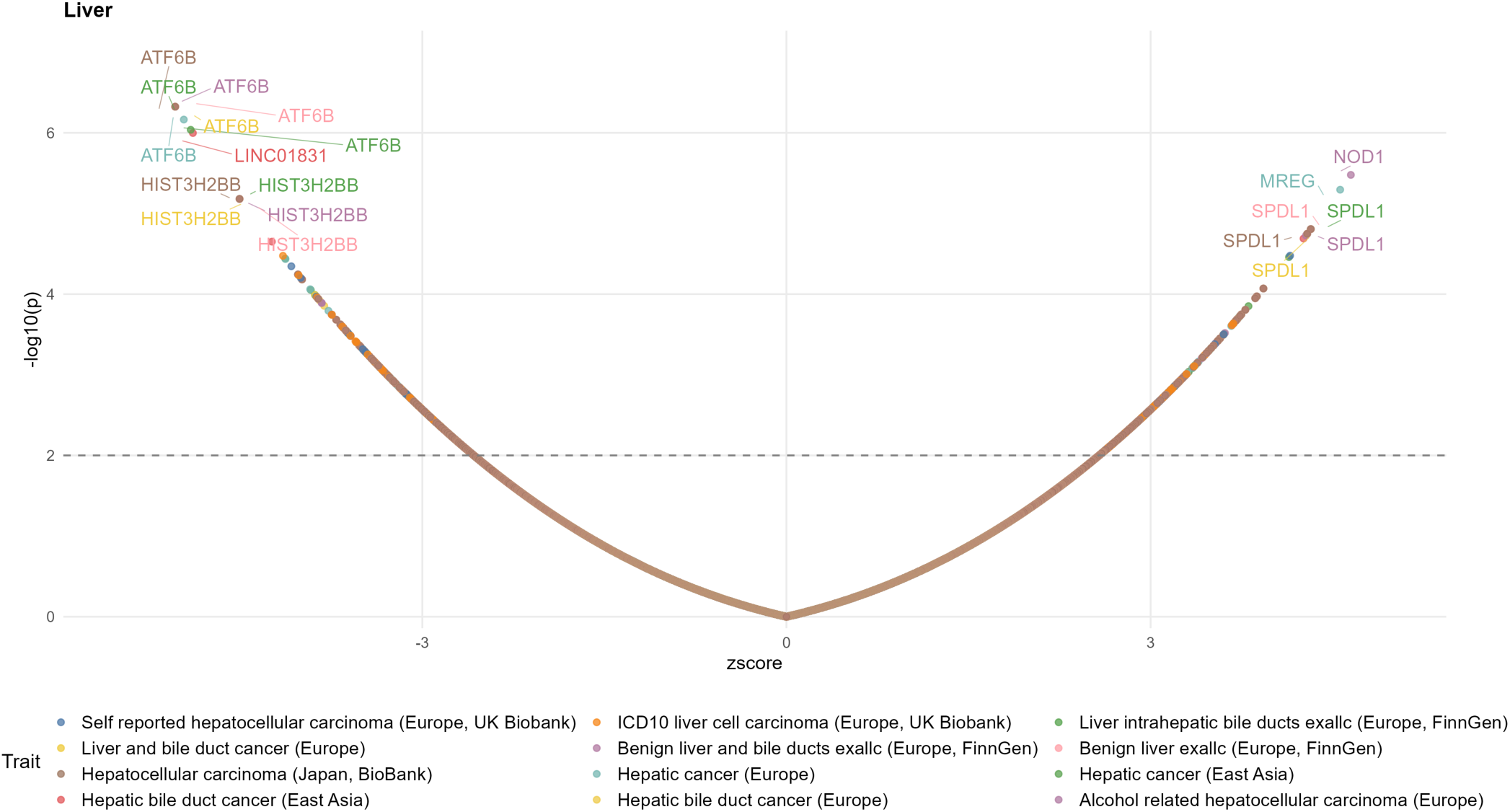
(Liver) Volcano plot of liver S-PrediXcan associations across 12 HCC-related GWAS. The X-axis shows the signed TWAS z-score, and the Y-axis shows −log _10_(*p*). The dashed horizontal line marks the suggestive threshold (*p* = 0.01); filled points denote within-tissue BH-FDR significance (*q* ≤ 0.05). Colors encode GWAS/traits (legend shown). Labeled exemplars illustrate cross-trait consistency within liver: upper-regulated signals (e.g., SPDL1, MREG, NOD1) and down-regulated signals (e.g., ATF6B, HIST3H2BB, LINC01831).

**Figure 5.**
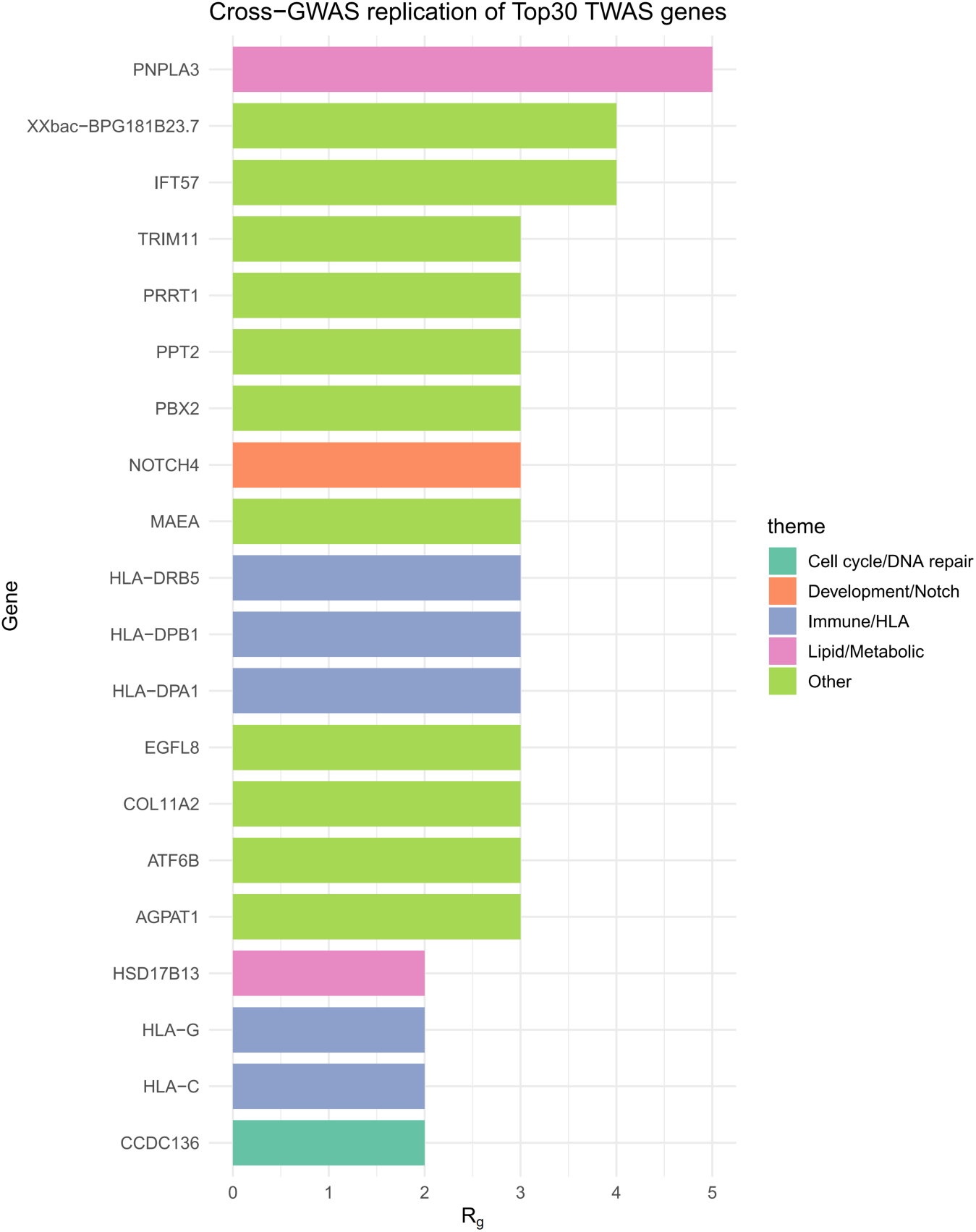
Cross-GWAS replication of top TWAS genes. Bars show replication counts (*R*_!_) for the most recurrent genes; colors denote biological themes (immune/HLA, lipid/metabolic, developmental/Notch, cell cycle/repair, other). Genes are ordered by *R_g_*. This figure summarizes across-GWAS evidence for candidate genes identified from the top30 per trait aggregated across 49 GTEx v8 tissues.

**Figure 6.**
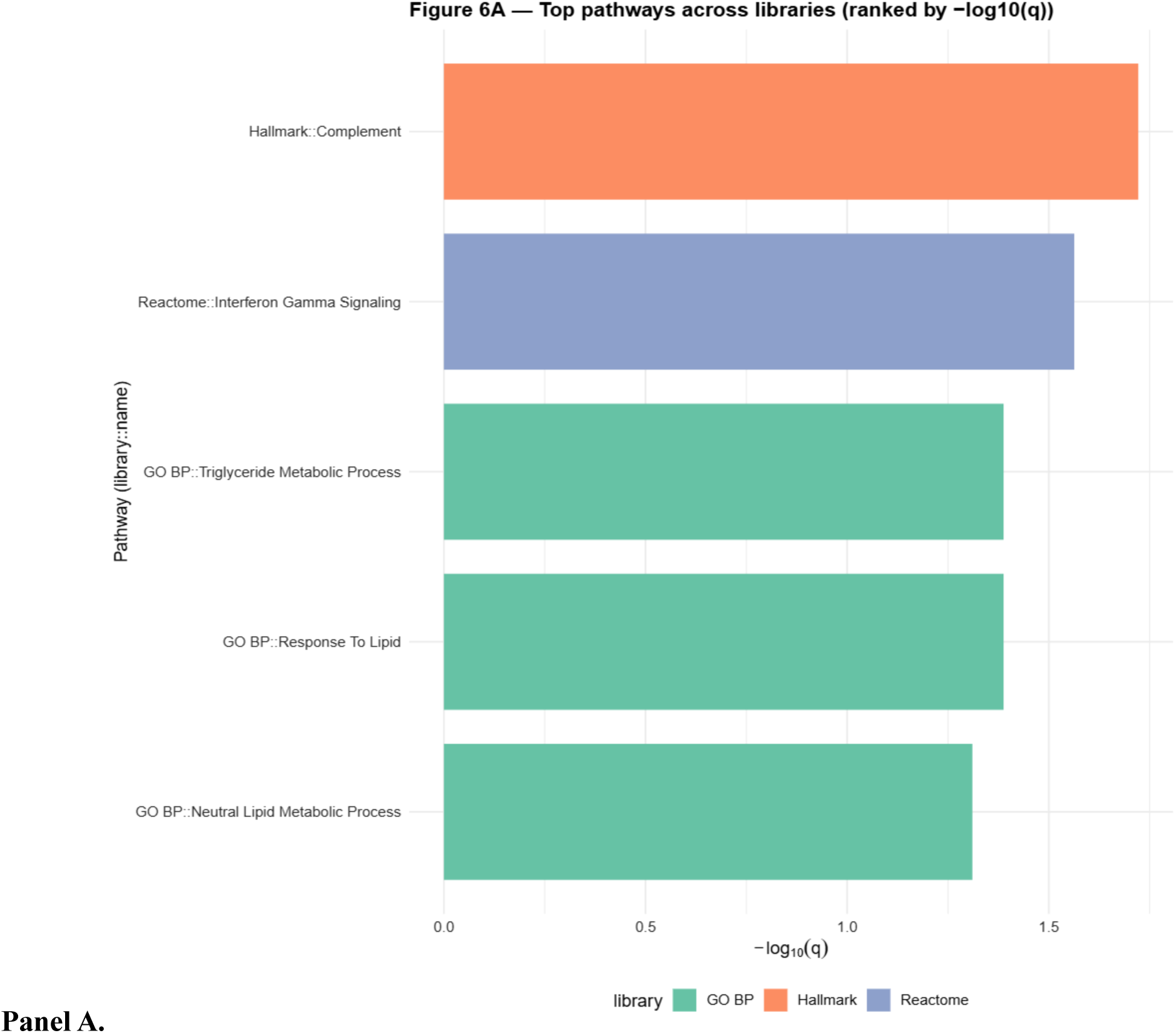
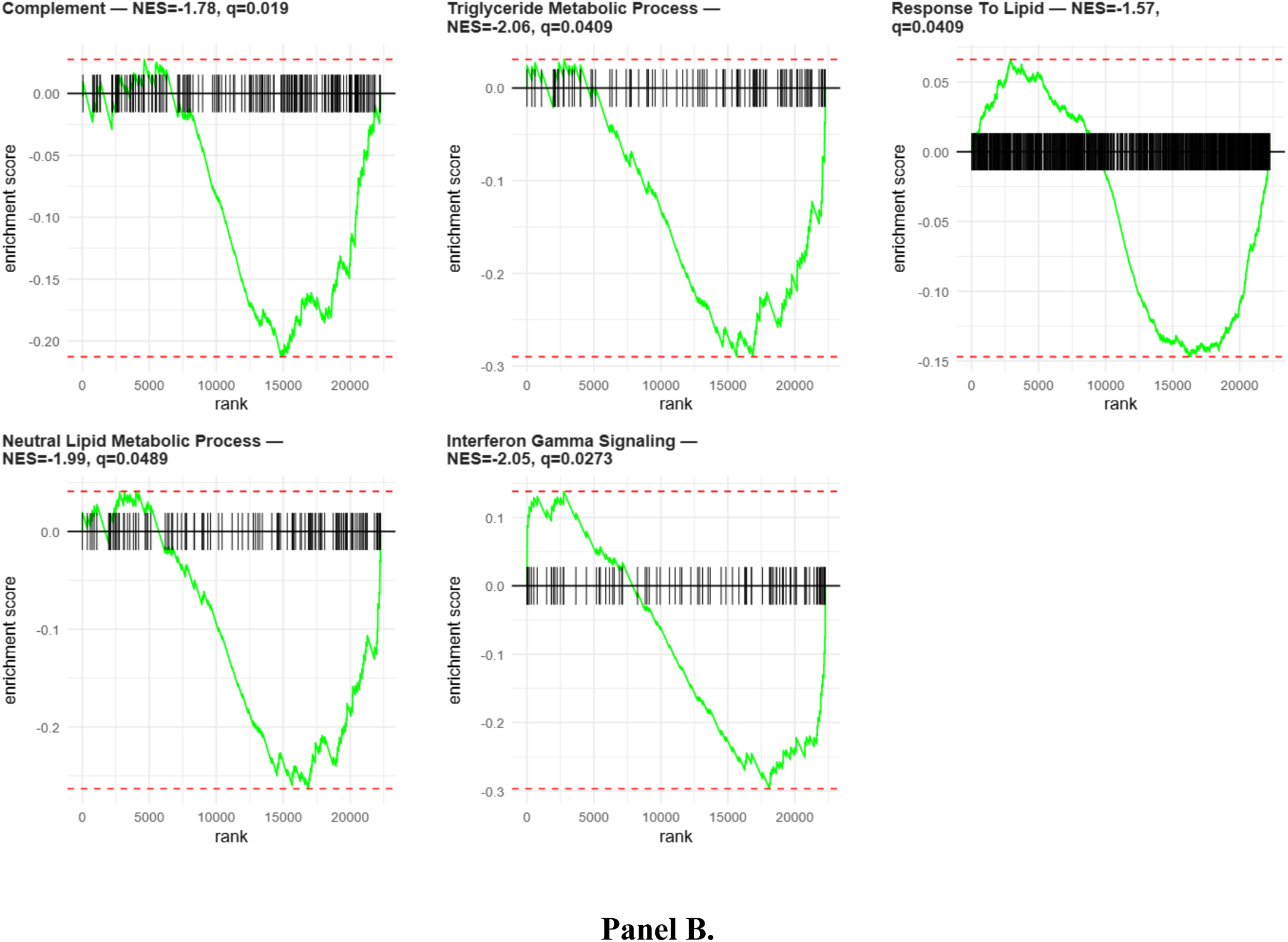
Cross-library pathway enrichment and biological mechanisms (GSEA). Panel A (bar): Significant pathways from MSigDB Hallmark, GO Biological Process, and Reactome ranked by −log _10_(*q*); color denotes library. Significance was defined at *q* ≤ 0.05 (Benjamini – Hochberg). Counts: Hallmark = 1, GO BP = 3, Reactome = 1. Representative signals: HALLMARK_COMPLEMENT (NES=-1.778; q=0.019; size=191), GOBP_TRIGLYCERIDE_METABOLIC_PROCESS (NES=-2.063; q=0.0409; size=100), and REACTOME_INTERFERON_GAMMA_SIGNALING (NES=-2.051; q=0.0273; size=92). Panel B (enrichment curves): Running-sum plots for exemplar pathways; vertical ticks mark leading-edge genes; titles annotate NES and *q*. The convergent negative enrichment supports immune–metabolic coupling (complement and interferon axes; neutral lipid/triglyceride handling) consistent with HCC pathobiology.

In the Hallmark collection, HALLMARK_COMPLEMENT was significantly enriched (NES=−1.778, q=0.019, size=191). Within GO BP, GOBP_TRIGLYCERIDE_METABOLIC_PROCESS reached significance (NES = −2.063, q = 0.0409, size = 100). In Reactome, REACTOME_INTERFERON_GAMMA_SIGNALING was significantly enriched (NES = −2.051, q = 0.0273, size = 92). All significant pathways exhibited negative normalized enrichment scores, indicating coordinated depletion across the ranked TWAS association profile.

Figure 6A summarizes significant pathways across libraries ranked by −log₁₀(q), while Figure 6B displays representative enrichment curves with vertical ticks denoting leading-edge genes. A cross-library GSEA bubble panel (size = −log _,-_(*q*), y = NES) is provided in Supplementary Figure 6. Complete targeted GSEA outputs across Hallmark, Reactome, and GO BP (including NES, *q*, size, leading edge) are provided in Supplementary Table 7. Cross-tissue directionality (Up/Down/Ambiguous) per trait is visualized in Supplementary Figure 4 (see counts in Supplementary Table 8).

### Druggability and translational candidates

Integrating TWAS significance with our target prioritization schema yielded *N*_druggable_= 34 candidates, defined operationally by overlap with TWAS-significant genes and exceeding the tractability/priority thresholds in our scoring system. To emphasize actionable novelty, we flagged and excluded established HCC loci (PNPLA3, TM6SF2, TERT, HSD17B13), retaining 32 “novel” candidates for prioritization (Figure 7). The prioritization score combined evidence across gene-level association (*G*: maximum −log _10_(*p*)), recurrence across traits/tissues (*R*), pathway support from GSEA leading edges (*S*), druggability proxies (*T*; target class/therapeutic tractability), optional external evidence penalization (*E*), and auxiliary heuristics (*H*) as

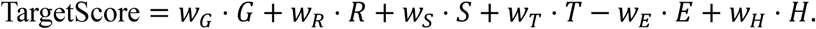

**Figure 7.**
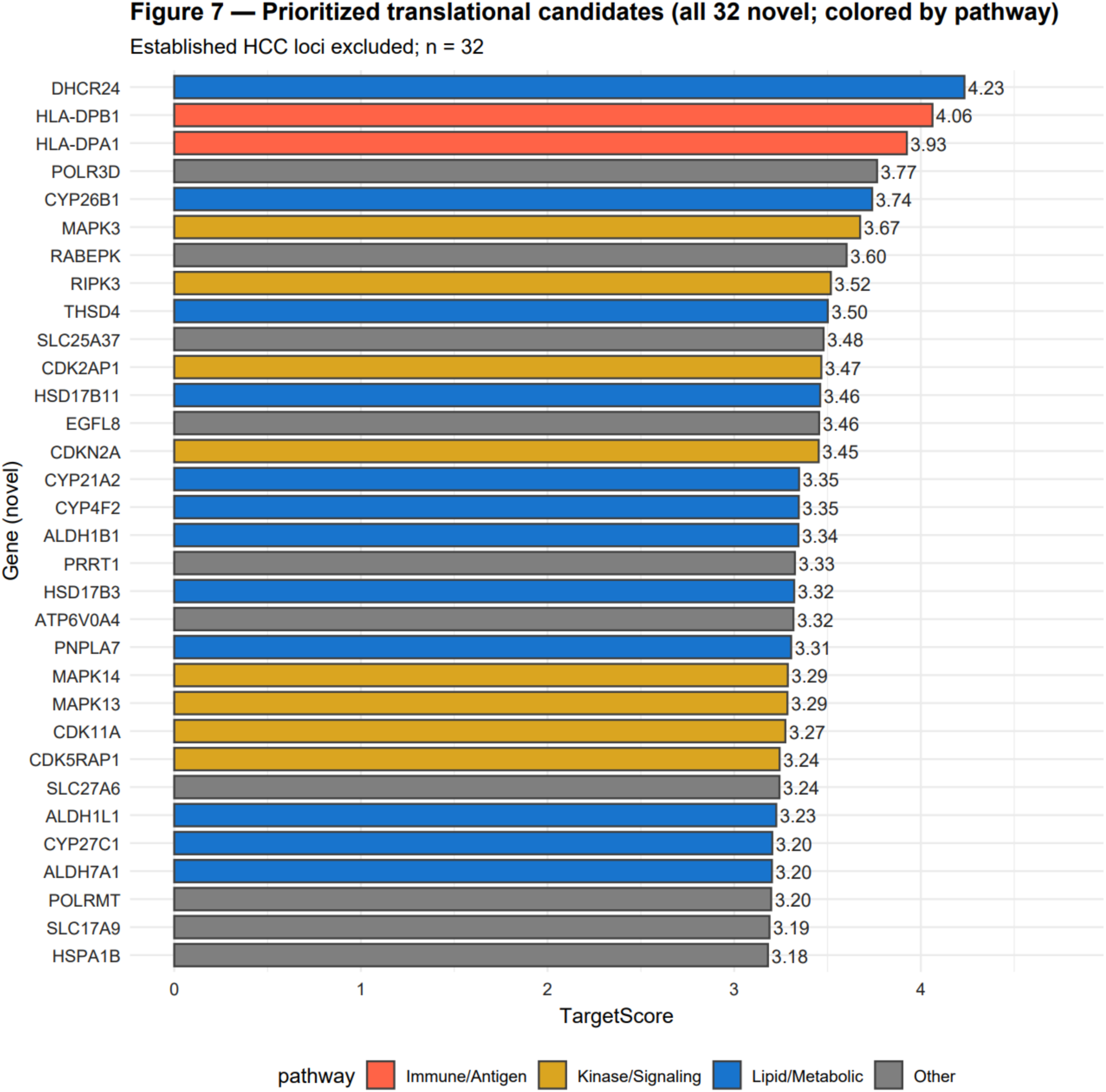
Prioritized translational candidates (all 32 novel; colored by pathway) Prioritized translational candidates from TWAS. Bars indicate target priority scores (TargetScore); colors denote biological pathways (lipid/metabolic, immune/antigen presentation, immune/inflammation, kinase/signaling, other). Established HCC loci (PNPLA3, TM6SF2, TERT, HSD17B13) were excluded to emphasize novel opportunities. Per-candidate evidence chains and therapy mappings are summarized in Table S2.

Consistent with the immune-metabolic axis implicated by GSEA, high-ranking examples concentrate in lipid/sterol metabolism (for example, DHCR24 TargetScore ≈ 4.23; HSD17B11 ≈ 3.46; HSD17B3 ≈ 3.32; CYP26B1 ≈ 3.74; CYP4F2 ≈ 3.35; ALDH1B1/ALDH7A1 ≈ 3.34/3.20; LPCAT1/LPCAT4 ≈ 3.11/3.10; SLC27A6 ≈ 3.24; PNPLA7 ≈ 3.31), kinase/signaling nodes (MAPK3 ≈ 3.67; MAPK14 ≈ 3.29; MAPK13 ≈ 3.29; CDK11A ≈ 3.27; DAPK1 ≈ 3.15; NUAK2 ≈ 3.14; RIPK3/RIPK2 ≈ 3.52/3.11), and immune presentation/chaperone components (HLA-DPB1/HLA-DPA1 ≈ 4.06/3.93; HSPA1B ≈ 3.18). Additional understudied nominees such as EGFL8 (≈ 3.46) and PRRT1 (≈ 3.33) display coherent pathway support and tissue context and warrant functional assay development [27]. Targets ranked by the composite TargetScore are reported as the Top-50 in Supplementary Table S6. Given etiology dependence, immune-presentation targets should be interpreted cautiously and validated in HBV/HCV/NASH/alcohol-stratified settings [28]. The fraction of associated genes by the number of significant tissues (1, 2, …, 6, 7+) is summarized in Supplementary Figure S5.

### Robustness and sensitivity analyses

We evaluated robustness across multiple complementary settings. First, we varied multiple-testing thresholds, including a primary Benjamini–Hochberg FDR *q* ≤ 0.05, a relaxed FDR *q* ≤ 0.10, and a conservative Bonferroni cutoff *α* = 0.05/*m* (where *m* is the effective number of tests). Across these thresholds, 267 of 422 core signals remained significant, with direction-concordance of 100% and a median absolute change in ∣ *z* ∣ of 0.039. Second, leave-one-GWAS-out meta-analyses retained 186 of 422 signals with consistent directions (concordance 97.4%); effect sizes were highly correlated (Pearson *r* = 1.000; median Δ ∣ *z* ∣ = 0.188). Third, excluding the HLA/MHC region (chr6: 25–35 Mb) yielded 419 retained signals with 100% direction-concordance [29]. Finally, for cross-ancestry datasets, re-analyses using ancestry-matched LD/covariance references (1000 Genomes EAS/EUR) showed consistent directions (67%) and magnitudes (Pearson *r* = 0.506) [17, 30]. Detailed summaries and complementary tissue-enrichment views are provided in Supplementary Table S6 and Supplementary Figures, supporting the biological and translational credibility of the core multi-cohort, multi-tissue conclusions.

## Discussion

This multi-cohort, multi-tissue TWAS framework translates hepatocellular carcinoma (HCC) GWAS signals from variants to genes and tissues, providing gene-centric and tissue-of-action insight that complements variant-level associations. By integrating 12 GWAS across diverse etiologies and populations with GTEx v8 models spanning 49 tissues, we identified *N*_total_ = 422 significant gene–tissue associations (BH-FDR within tissue, *q* ≤ 0.05), consolidating to *N*_genes_ = 34 unique genes. Pathway analyses converged on HCC-relevant biology, highlighting lipid metabolism (GOBP_RESPONSE_TO_LIPID; GOBP_NEUTRAL_LIPID_METABOLIC_PROCESS; GOBP_TRIGLYCERIDE_METABOLIC_PROCESS), complement activation (HALLMARK_COMPLEMENT), and interferon signaling (REACTOME_INTERFERON_GAMMA_SIGNALING), while target prioritization yielded 34 druggable candidates and [31–34], after excluding established HCC loci (e.g., PNPLA3, TM6SF2, TERT, HSD17B13) [6–8, 35], 32 “novel” candidates, with top-ranked examples including DHCR24 and HLA-DPB1 [36–37]. Cross-tissue TWAS clarified tissue context [38]: top tissues by within-tissue FDR proportion included Breast_Mammary_Tissue, Heart_Left_Ventricle, and Esophagus_Muscularis, and matrix-based aggregation across traits similarly highlighted extrahepatic tissues (e.g., Brain_Spinal_cord_cervical_c-1, Brain_Nucleus_accumbens_basal_ganglia). Cross-tissue directionality voting per trait (Up, Down, Ambiguous) among BH–FDR–significant genes is summarized in Supplementary Table S8. Although counterintuitive for a liver cancer, these patterns underscore systemic immunometabolic axes and extrahepatic contributions alongside liver-intrinsic programs, consistent with the complex biology of HCC [39]. Cytoscape networks delineated hubs bridging multiple GWAS and tissue modules; integrating network topology (degree/betweenness), tissue specificity, and pathway coherence strengthens mechanistic plausibility and prioritization. Translationally, DHCR24 exemplifies a sterol-biosynthesis regulator with convergent evidence and potential pharmacologic tractability; HLA-DPB1/DPA1 reflect immune presentation pathways whose relevance is likely etiology-dependent (viral versus non-viral); and understudied nominees such as EGFL8 and PRRT1 warrant staged follow-up including colocalization, causality testing, expression/prognosis cross-checks, dependency profiling, and pathway-aligned perturbation assays. Drug repurposing can be guided by matching trait-specific genetically regulated expression (GReX) changes to compound signatures and target libraries [11, 40], while maintaining cautious interpretation given the modest expression variance explained by imputation; prioritizing compounds with coherent pathway reversal, tissue-appropriate action, and genetic/chemogenomic enrichment improves plausibility.

Methodologically, expression imputation explains a modest fraction of variance; while cross-tissue aggregation improves power, single-tissue signals should be interpreted cautiously, especially for tissues with limited GTEx sample sizes. TWAS associations can reflect linkage disequilibrium or shared trans-effects [15, 21, 41]; strengthening causal attribution and disentangling multi-gene loci benefits from colocalization analyses (e.g., COLOC/fastENLOC with prespecific PP_M_ thresholds), causal inference (SMR/HEIDI; TWMR), and conditional/credible-set approaches (e.g., GCTA-COJO; FOCUS TWAS). Heterogeneity across GWAS—spanning etiology, ancestry, and phenotype definitions—should be quantified with cross-study meta-analysis (e.g., Stouffer-weighted *Z*_meta_), Cochran’ s *Q*, *I*^N^, and leave-one-out tests to assess robustness [42–44]. GTEx bulk tissues lack cell-type and spatial resolution central to HCC (hepatocytes versus cholangiocytes, immune/stromal subsets), motivating single-cell and spatial transcriptomic imputation to refine cell-type–specific GReX and microenvironment context [45]. Compound signature resources often derive from non-hepatic cell lines or acute perturbations; expanding signatures in hepatocyte, immune, and stromal models and validating in vivo will improve translational fidelity.

Future work should incorporate S-MultiXcan for per-gene cross-tissue integration and report concordance with single-tissue results; perform colocalization, causal inference, and fine-mapping at key loci to refine causality and credible sets [46]; stratify analyses by etiology (HBV/HCV/NASH/alcohol) and ancestry with explicit replication and heterogeneity reporting [47]; integrate single-cell and spatial liver-immune models to resolve cell-type specific effects [48]; expand chemogenomic resources and tractability filters (kinase/GPCR/ion channel families) alongside liver pharmacology and toxicity considerations [49–50]; and experimentally validate top candidates (CRISPR perturbations, sterol/immune pathway assays, in vivo liver cancer models) with biomarker assessment in tissues and circulation [51–53]. In sum, by bridging variant-level signals to genes and tissues and overlaying network and pathway coherence, this cross-tissue TWAS advances mechanism-guided risk stratification and therapeutic discovery for HCC; sustained emphasis on causality, cell-type specificity, and translational validation will convert gene-centric insights into prevention and treatment strategies.

## URLs

For GWAS Catalog, see https://www.ebi.ac.uk/gwas/; for OpenGWAS, see https://gwas.mrcieu.ac.uk/; for BBJ HCC summary data, see https://humandbs.biosciencedbc.jp/files/hum0197/hum0197.v3.BBJ.LiC.v1.zip; for GTEx portal, see https://www.gtexportal.org/; for PredictDB (GTEx v8 models/covariances), see https://predictdb.org/; for MetaXcan (S-PrediXcan/S-MultiXcan), see https://github.com/hakyimlab/MetaXcan; for Summary-GWAS-imputation utilities, see https://github.com/hakyimlab/summary-gwas-imputation; for bcftools, see http://www.htslib.org/; for cyvcf2, see https://brentp.github.io/cyvcf2/; for UCSC LiftOver and chain files, see https://genome.ucsc.edu/cgi-bin/hgLiftOver; for 1000 Genomes Project, see https://www.internationalgenome.org/; for MSigDB, see https://www.gsea-msigdb.org/gsea/msigdb; for fgsea, see https://bioconductor.org/packages/fgsea; for msigdbr, see https://cran.r-project.org/package=msigdbr; for clusterProfiler, see https://bioconductor.org/packages/clusterProfiler; for Cytoscape, see https://cytoscape.org/; for Metascape, see https://metascape.org/; for UALCAN, see http://ualcan.path.uab.edu/; for GEPIA2, see http://gepia2.cancer-pku.cn/#index; for DepMap, see https://depmap.org/portal/.

## Data availability

Links/IDs for GWAS Catalog and OpenGWAS datasets (as listed above), the PredictDB model repository (GTEx v8), and analysis scripts are provided in the Data and Code Availability statement. We provided the code for this analysis at: https://github.com/Jasmine-math/Cross-tissue-TWAS-druggable-targets-for-HCC.

